# Changes in surface temperatures reveal the thermal challenge associated with catastrophic moult in captive Gentoo penguins

**DOI:** 10.1101/2024.01.16.575878

**Authors:** Agnès Lewden, Tristan Halna du Fretay, Antoine Stier

**Affiliations:** Université de Brest - UMR 6539 CNRS/UBO/IRD/Ifremer, Laboratoire des sciences de l’environnement marin - IUEM - Rue Dumont D’Urville - 29280 - Plouzané; Université de Strasbourg, CNRS, IPHC UMR 7178, F-67000 Strasbourg, France; Department of Biology, University of Turku, Turku, Finland

**Keywords:** Thermal challenge - Moult, Thermoregulation, Penguin, Global warming

## Abstract

Once a year, penguins undergo a catastrophic moult replacing their entire plumage during a fasting period on land or on sea-ice during which time individuals can lose 45% of their body mass. In penguins, new feather synthesis precedes the loss of old feathers leading to an accumulation of two feathers layers (double coat) before the old plumage is shed. We hypothesize that the combination of the high metabolism required for new feathers synthesis and the potentially high thermal insulation linked to the double coat could lead to a thermal challenge requiring additional peripheral circulation to thermal windows to dissipate extra-heat. To test this hypothesis, we measured the surface temperature of different body regions of captive Gentoo penguins (*Pygoscelis papua*) throughout the moult under constant environmental conditions.

The surface temperature of the main body trunk decreased during the initial stages of the moult, therefore suggesting a higher thermal insulation. On the opposite, the periorbital region, a potential proxy of core temperature in birds, increased during these same early moulting stages. The surface temperature of bill, flipper and foot (thermal windows) tended to initially increase during the moult period, highlighting the likely need for extra heat dissipation in moulting penguins. These results raise questions regarding the thermoregulatory capacities of wild penguins during the challenging period of moulting on land in the current context of global warming.

## Introduction

In birds, feathers have many functions including flight, thermal insulation, communication (with plumage coloration; e.g. Bortolotti 2006) as well as tactile sensation (Cunnighams et al. 2011). Plumage provides thermal insulation for endothermic birds helping them to maintain a high core body temperature (Prinzinger et al. 1991). Indeed, the feather layers trap air above the skin (Dawson et al. 1999) and plumage color and microstructure of plumage elements (Wolf and Walsberg 2000) reduce conductive, convective and radiative heat loss between bird and the outside environment (e.g. Calder and King 1974; Bakken 1976; Wolf and Walsberg 2000). This is especially true in aquatic birds such as penguins that show a high density of downy and contour-feathers increasing the water resistance (Pap et al. 2017; Osváth et al. 2018). Penguins have a thick and morphologically specialized plumage (Rutschke 1965; Williams et al. 2015) providing 80-90% of insulation requirements (Le Maho et al. 1976; Le Maho 1977) that enable them to exist in the harshest climates of Antarctica. It is therefore important that penguins are able to maintain high quality plumage (Jenni and Winkler 2020), through the moult: a replacement of the old and damaged feathers by new ones (Humphrey and Parkes 1959).

The moult of penguins is described as “catastrophic” (Davis and Darby 2012) and occurs once a year during a fasting period on land or on sea-ice where the heat conductance of air is 25 times lower than that of water (de Vries and van Eerden 1995). During this time, individuals replace their entire plumage in two overlapping stages with the synthesis of new feathers preceding the loss of old feathers (Groscolas and Cherel 1992; Fig. 1A). New feathers begin to grow under the skin until they reach 40% of their size, when they emerge through the skin. Between 40% and 60% of the new feather growth, the old feathers remain attached to the new feathers and at this stage birds simultaneously have two feather layers (Fig. 1A). The old feathers then fall off, reducing thermal insulation until the new feathers finish growing (Groscolas and Cherel 1992; Fig. 1A). Moult is an energetically costly period for penguins (Croxall 1982; Adams and Brown 1990), despite their low level of activity while fasting on land (Cherel et al. 1994). Indeed, the metabolic rate increases by a factor of 1.3 and 1.5 in king penguins (*Aptenodytes patagonicus*; Cherel et al. 1994) and in little penguins (*Eudyptula minor*; Baudinette et al. 1986) respectively. During this fasting period, macaroni (*Eudyptes chrysolophus*) and rockhopper penguins (*E. chrysocome*) lose 44% and 45% of their body mass respectively during a 25-day moult period (Brown 1986). Similarly king and emperor penguins (*Aptenodytes forsteri*) lose approximately 45% of their body mass in 30 days with a peak of daily body mass loss during the final stage of feather loss (Groscolas 1978; Cherel et al. 1988).

**Fig. 1:**
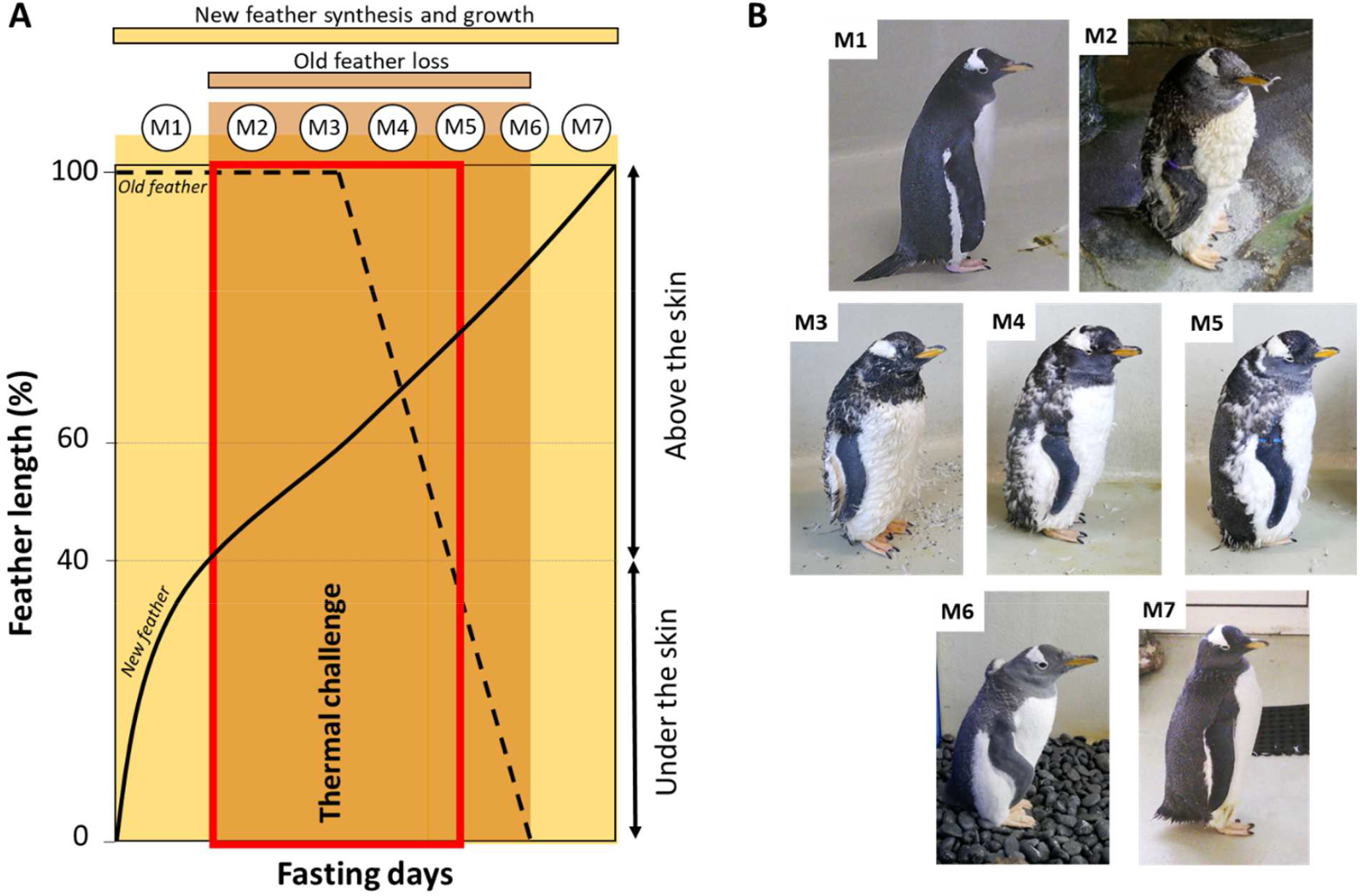
Identification of the seven main moulting stages (stages M1 to M7) characterized during the Gentoo moult in captivity. The schematic representation of the penguin moult **(A)** shows the new feathers (straight-line) grow beneath the outer skin layer until 40% of their total length. Between 40 and 60% of new feather growth, penguins have a double feather layer, old (dashed line) and new. These two feather layers could lead to a thermal challenge for heat dissipation (red box). After 60% of new feather growth, the old feathers are starting to fall-off. Adapted from Groscolas and Cherel (1992). During moult, visual plumages changes can be noticed **(B)** and characterized into seven different stages: M1 = uniform old plumage, M2 = ‘pop-corn’ (superposition of old and new immature feathers), M3 = first fall of old plumage, M4 = 25% of old feathers fallen, M5 = 50% of old feathers fallen, M6 = 75% of old feathers fallen and M7 = uniform new plumage corresponding at the end of the monitoring. See the main text for more details ©A. Lewden

While Groscolas and Cherel (1992) suggested a decrease in thermal insulation during the loss of old feathers, the preceding overlay of the new and the old feathers could increase the overall thermal insulation of plumage. Metabolic heat production increases during the moult due to feather synthesis and increased peripheral blood flow to grow the feather. During the earlier stages of the moult, when penguins may have a greater thermal insulation due to the double layer of feathers, penguins may face a thermal challenge (Fig. 1A) by being unable to efficiently dissipate metabolic heat, potentially leading to a rise in body temperature. To investigate this hypothesis, we measured surface temperatures of captive Gentoo penguins (*Pygoscelis papua*) using thermal imaging during the entire moulting period. The captive conditions allowed us to measure individuals throughout the full moult period at a uniform air temperature without interferences of solar radiation, wind or precipitations. We measured surface temperature of old and new plumage to represent well-insulated body regions, periorbital region as a proxy of core temperature (e.g. Gauchet et al. 2022), while surface temperature of bill, flipper and the foot correspond to thermal windows (i.e. poorly insulated body areas governed by a vascular system that controls their surface temperature; Tattersall et al. 2009; McCafferty et al. 2013; Lewden et al. 2020). Specifically, we predicted that when penguins possessed two simultaneous feather layers (moulting stages M2 to M5; Fig. 1) there would be a decrease of plumage surface temperature, an increase in surface temperatures of thermal windows and a potential rise in the temperature of the periorbital region.

## Material and Methods

### Study site

Twenty-seven Gentoo penguins were studied in captivity at Océanopolis© aquarium, Brest, France. Individuals were identified by a colored plastic ring on the right flipper, and divided into two groups, the first of 18 individuals (7 males and 11 females) and the second of 13 individuals (6 males and 7 females). Individuals were maintained within their thermoneutral zone, that is between 8 and 15°C in Gentoo penguins (Taylor 1985; Wilson et al. 1998), in two separate enclosures with similar conditions, *i*.*e*. a permanent access to free water, unfed during moulting period and with the same number of enclosure cleaning and animal keeper visits. During measurement sessions, air temperature (*T*_a_) and relative humidity (RH) were measured using a weather station Kestrel® 5400 Heat Stress Tracker. The wet-bulb temperature (*T*_w_) was then calculated according to the equation (1) in Stull 2011, to take into account the cooling effect of higher humidity. Enclosures showed a relatively stable *T*_w_ during the study period (from July, 30^th^ to October, 20^th^) with a range of temperature between 7.20 and 12.56°C in the first group (Group 1) and between 9.19°C and 14.20°C in the second group (Group 2). However, we measured a small but significant difference between groups/enclosures with a higher *T*_w_ in Group 2 (mean ± standard error of 10.82 ± 0.39°C) compared to Group 1 (mean ± standard error of 9.38 ± 0.28°C) (P<0.005). Similarly, the ground surface temperature (*T*_ground_) in contact with penguin’s feet was higher in Group 2 (mean ± standard error of 14.65 ± 0.13°C) compared to Group 1 (mean ± standard error of 12.93 ± 0.09°C)(P<0.0001). Moult lasted 14.0 ± 0.66 days per individual in Group 1 and 12.8 ± 0.97 days per individual in Group 2, without significant difference between group (P=0.85).

### Moult

Penguin surface temperatures were measured once a day in the morning, at least twice to a maximum of 25 times per individuals with a mean of 11.75 measurements per individual. To track the progress of the moult, 7 moult stages were characterized (Fig. 1B) ranging from a uniform old-plumage (Fig. 1B; M1) to a uniform new-plumage (Fig. 1B; M7) and assigned by the same observer (A.L) during data collection. The intermediate stages were “pop-corn” during which individuals carried two feather layers giving them a puffy appearance (Fig. 1B; M2). The “first fall” stage corresponds to the advanced pop-corn stage with the first fall of old feathers visible (Fig. 1B; M3). The “25%” stage corresponds to the acceleration of old feather fall, with at least 25% of the trunk having lost its old plumage and thus presenting a new, immature plumage (Fig. 1B; M4). The “50%” stage corresponds to the peak of the moult, with 50% of the trunk showing two layers of plumage and 50% showing new immature plumage (Fig. 1B; M5). At the “75%” stage (Fig. 1B; M6), individuals had lost most of their old plumage and the new plumage, while not yet fully grown, visibly increased in volume.

### Thermal image collection and analysis

One or two thermal pictures of left or/and right profiles were taken per day of measurement from the same angle (Playà-Montmany and Tattersall 2021) at a distance of *ca*. 1 meter with each body area being larger than ten times the spot size (Tabh et al. 2021) of 0.65 mm from our FLIR E96 thermal camera (640×480 pixel). For each bird, profile pictures were defined as right (ringed side) or left (non-ringed) side. We hypothesized that the ringed flipper might show a higher surface temperature induced by a light inflammation linked to ring friction. Emissivity was set to 0.98 (Whittow 1986; Monteih and Unsworth 1990) wheareas *T*_a_ and RH were set for each picture using Flir ThermaCAM Researcher Professional 2.10 software. Bill, flipper and foot areas were delineated by tracing a polygon around the edge to extract the mean surface temperature of each area (hereafter *T*_bill_, *T*_flipper_, *T*_foot_ respectively). Mean surface temperature of the ground (*T*_ground_) was extracted using a standard square size (877 pixel²) situated just below the feet. Head was also delineated and the maximum surface temperature of this area was extracted corresponding to the periorbital region (hereafter *T*_eye_; Jerem et al. 2018). As the loss of old plumage on the trunk was not symmetrical on both sides, we could not determine a representative trunk surface temperature from the profile pictures. This is why, we calculated an index of (hereafter *T*_trunk_) using the old and the new plumage surface temperature (*T*_old plumage_ and *T*_new plumage_ respectively) according to the percentage of trunk recovery as following:

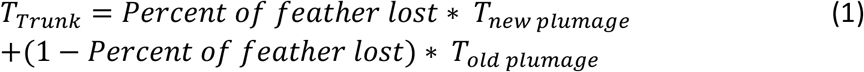

*T*_old plumage_ and *T*_new plumage_ corresponded to a mean surface temperature of a standard square size (438 pixel²) positioned on a uniform patch of each kind of plumage. When individuals were in old plumage (M1) in pop-corn (M2) and in first fall (M3) stages, *T*_trunk_ corresponded to *T*_old plumage_. When individuals were in new plumage stage (M7) *T*_trunk_ corresponded to *T*_new plumage_.

For others moulting stages (M4; M5 and M6), *T*_trunk_ was calculated using the equation (1) determined from the cover of the entire trunk circumference to weight the proportion of each kind of plumage.

### Statistical analysis

The effect of plumage color (black or white) on old and new plumage surface temperatures was initially investigated using an ANOVA, but dropped out of final models since as expected in the absence of solar radiation, we did not find any significant difference in surface temperature between black and white plumage in both old and new plumage (*P*>0.80 in both cases). Similarly, we did not measure any significant effect of the ring on *T*_flipper_ and therefore excluded this variable from our final statistical models (*P*=0.99).

The relationship between four temperature areas (*i*.*e*. bill, periorbital, flipper and trunk) and the moult stages were studied using general linear mixed models (GLMM), including penguin ID as a random intercept, to control for repeated measures and *T*_w_, Group (1 or 2) and Sex (male or female) as fixed effects. For the *T*_foot_, *T*_ground_ replaced *T*_w_ in the model considering the large surface of foot in contact with the ground (*i*.*e*. conductive heat loss). Indeed, the linear relationship between *T*_w_ and *T*_foot_ (R²= 0.02; P=0.004) was markedly weaker than the relationship between *T*_ground_ and *T*_foot_ (R²=0.38; P<0.0001).

Differences were then investigated using a Tukey’s honestly significant difference (HSD) *post-hoc test* for moult stages effect. Statistical analyses were performed using JMP® v. 13 (SAS Institute Inc., Cary, North Carolina, USA) and results were reported as mean ± standard error unless specified.

## Results

Body surface temperatures were positively related to *T*_w_ or *T*_ground_ (Table 1). The variable ‘group’ was only significant for *T*_trunk_ (*P*<0.0001) with lower *T*_trunk_ measured in the group exposed to the slightly colder environment (*i*.*e*. Group 1). Females had a slightly higher *T*_eye_ than males (31.02 ± 0.11 and 30.71 ± 0.10°C respectively; *P*=0.023). Group and sex did not explain any significant variation in *T*_bill_, *T*_flipper_ and *T*_foot_ (Table 1). However, variation in all body surface temperatures were significantly influenced by the stage of moult (Table 1).

**Table 1:** Summary of the general linear mixed models used to investigate variation in body surface temperatures during moult in captive Gentoo penguins.

| | | $T_{\text{eye}}$ | | | $T_{\text{trunk}}$ | | | $T_{\text{bill}}$ | | | $T_{\text{flipper}}$ | | | $T_{\text{foot}}$ | | |
| --- | --- | --- | --- | --- | --- | --- | --- | --- | --- | --- | --- | --- | --- | --- | --- | --- |
| | | $R^2=0.55$ N= 339 | | | $R^2=0.68$ N= 340 | | | $R^2= 0.36$ N= 340 | | | $R^2=0.31$ N= 340 | | | $R^2=0.60$ N= 327 | | |
| Random effect : |  | Variance |  |  | Variance |  |  | Variance |  |  | Variance |  |  | Variance |  |  |
| Bird ID |  | 0.05 |  |  | 0.29 |  |  | 0.25 |  |  | 0.16 |  |  | 3.49 |  |  |
| Residual |  | 0.91 |  |  | 1.18 |  |  | 9.38 |  |  | 6.05 |  |  | 14.03 |  |  |
| Fixed effect : |  | df | F | P | df | F | P | df | F | P | df | F | P | df | F | P |
| Ta or Tground |  | 1,328 | 6.05 | 0.014 | 1,320 | 42.39 | <.0001 | 1,330 | 20.81 | <.0001 | 1,330 | 18.66 | <.0001 | 1,316 | 172.19 | <.0001 |
| Group |  | 1,64 | 0.75 | 0.38 | 1,103 | 104.33 | <.0001 | 1,84 | 0.10 | 0.75 | 1,55 | 1.61 | 0.21 | 1,107 | 2.59 | 0.11 |
| Sex |  | 1,37 | 5.59 | 0.023 | 1,71 | 1.26 | 0.27 | 1,48 | 3.48 | 0.068 | 1,30 | 1.07 | 0.31 | 1,70 | 1.76 | 0.19 |
| Molt stages |  | 6,323 | 49.67 | <.0001 | 6,321 | 4.94 | <.0001 | 6,324 | 16.47 | <.0001 | 6,319 | 15.23 | <.0001 | 6,309 | 5.29 | <.0001 |

*T*_eye_ significantly increased from the old plumage to the pop-corn stage, and stayed elevated until the 50% moult stage (Fig. 2A). *T*_eye_ then decreased back at 75% of the moult (at a level similar to the old plumage), and ended up being the lowest at the new plumage stage (Fig. 2A). *T*_trunk_ showed an initial drop from the old plumage stage to the First fall stage (Fig. 3), and then increased back to its initial level as the moult progressed further towards the new plumage stage (Fig. 2A). *T*_trunk_ at the new plumage stage (15.93 ± 0.14°C) did not significantly differ from the old plumage stage (16.48 ± 0.21°C; *P*=0.15) (Fig. 2A and 3).

**Fig. 2:**
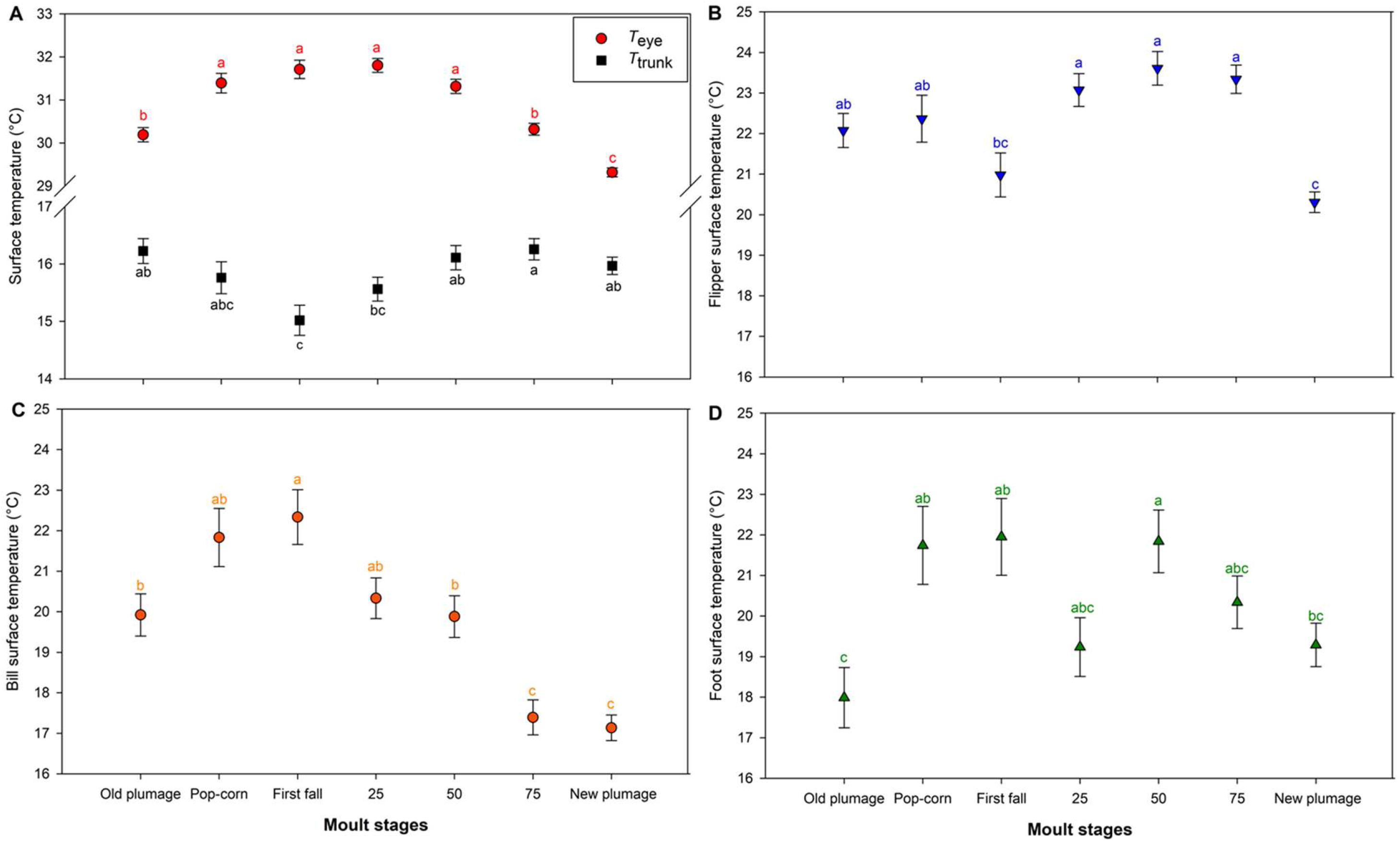
Least-squared mean surface temperatures during the seven moulting stages. Values for the periorbital region (red) and the trunk (black) (A), the flipper (B), the bill (C), and the foot (D) are shown. Values that do not share the same letter are significantly different from each other (post-hoc Tukey’s HSD test; P<0.05).

**Fig. 3:**
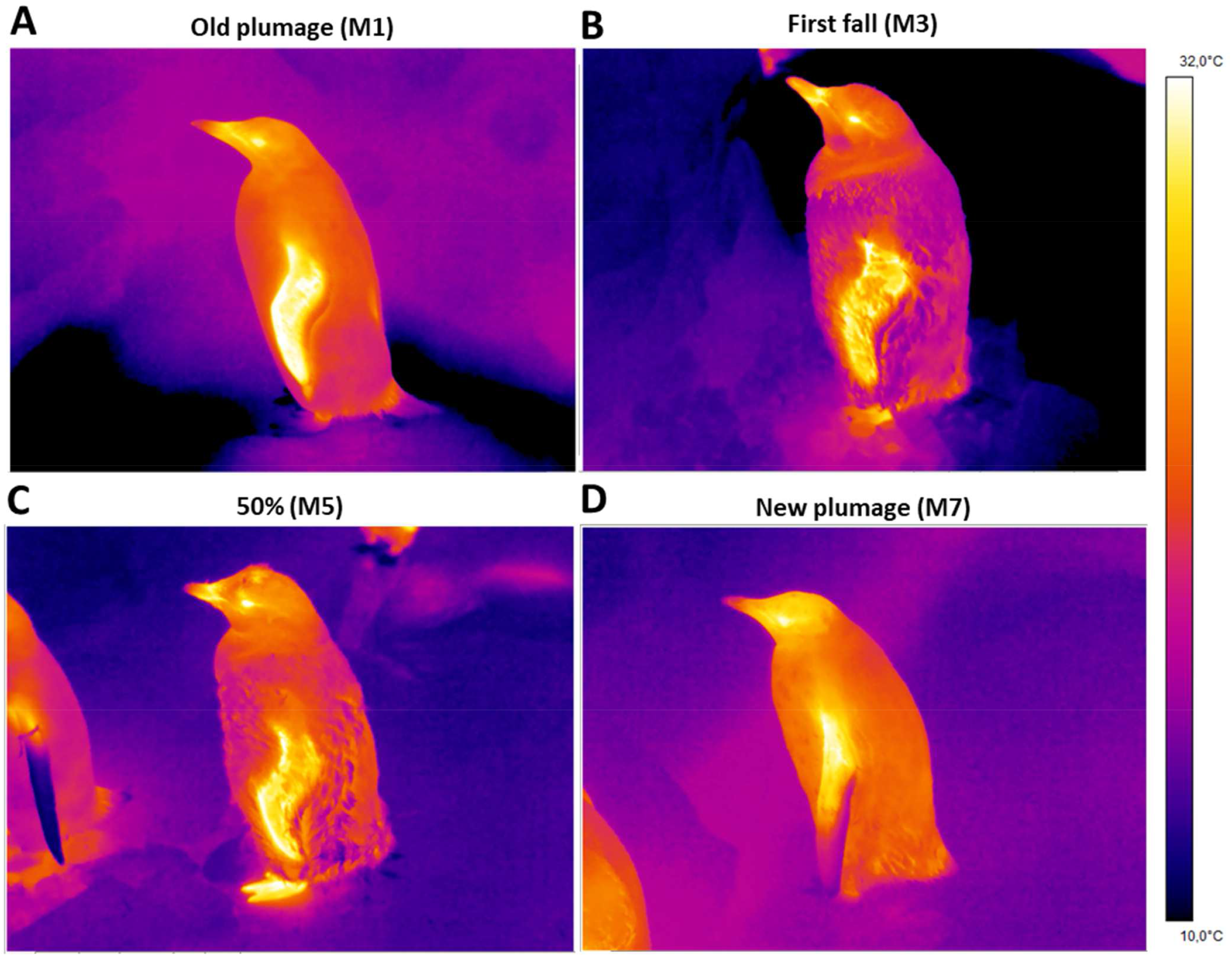
Visual comparison of four Gentoo penguins measured during moult by thermal imaging. Individuals were measured through the moult including before moult in old plumage **(A)**, at moulting stage first fall **(B)** and 50% **(C)** and in new plumage **(D)**. Images illustrate lower *T*_trunk_ at first fall (M3), lower *T*_flipper_ in new plumage (M7) compared to others stages and higher, higher *T*_bill_ at first fall (M3) than in new plumage (M7) and higher *T*_foot_ during moult (M3 and M5) than before and at the last stage of the moult (M1 and M7).

*T*_flipper_ had the highest temperature at stage 25, 50 and 75% of the moult (Fig. 2B). *T*_flipper_ at those stages were higher than at first fall and new plumage stages (all *P*<0.04; Fig. 3), but did not significantly differ from the old plumage and pop-corn stages (all *P*>0.30). *T*_flipper_ was 1.77°C colder at new plumage stage compared to old plumage stage (P=0.0002; Fig. 3).

*T*_bill_ significantly increased from the old plumage to first fall stage, and decreased thereafter (Fig. 2C) with a maximum difference of -5.20°C between the first fall and the new plumage stages (*P*<0.0001; Fig. 3). *T*_bill_ was also lower at the new plumage stage (17.13 ± 0.31°C) than at the old plumage stage (19.92 ± 0.52°C; P<0.0001).

*T*_foot_ initially increased from the old plumage until the First fall stage (Fig. 3), and remained slightly elevated until the end of the monitoring (i.e. new plumage; Fig. 2D). Yet, there was no significant difference in *T*_foot_ between new plumage (19.29 ± 0.53°C) compared to old plumage stages (17.99 ± 0.74°C; P=0.62).

## Discussion

Our study investigated the effect of moulting in captive Gentoo penguins, a fasting period during which metabolic rate increases (Baudinette et al. 1986; Cherel et al. 1994) while body insulation is heavily modified in penguin species (Groscolas and Cherel 1992). Our results showed that at early stages of the moult, when individuals have two feather layers (Stages M2, M3 and M4; Fig. 1 and 2), plumage insulation was elevated as shown by the lowest *T*_trunk_ (Fig. 2 and 3), while the surface temperatures of thermal windows (bill, flipper and foot) and the periorbital region were generally elevated at these stages (Fig. 2). This effect was maintained until approximately 50% percent of the moult, after which surface temperatures of non-insulated body regions started to decrease (Fig. 2 and 3), such that the mean surface temperatures of new plumage stage were significantly lower (except for *T*_*foot*_) than at old plumage stage (Fig. 2).

Early moulting stages in penguins may therefore provide a thermal challenge for birds to dissipate extra-heat. Moult is energetically costly (Baudinette et al. 1986; Cherel et al. 1994) through maintaining a peripheral blood flow for dermal perfusion sustaining feather synthesis. Simultaneously with this higher heat production, we found that the potential for heat dissipation is likely to be reduced by the additional insulation resulting from the combination of newly growing and old feathers, as shown by the tendency of *T*_trunk_ to decrease at pop-corn (M2), first fall (M3) and 25% of old feathers fallen (M4) stages (Fig. 2). Correspondingly, uninsulated or less insulated body areas from the bill, flipper and foot exhibited the opposite pattern (Fig. 2 and 3) with an increase of surface temperature allowing greater heat loss through radiation and convection to the surroundings. Within our captive experimental conditions, birds were measured within their thermoneutral zone (between 8° and 15°C; Taylor 1985), which theoretically enable to exclude any changes in metabolic rate associated with thermoregulation. Thus, our study highlights that blood flow to thermal windows may help to compensate for greater insulation and heat production associated with feather synthesis, in order to help maintaining a stable core body temperature during early moulting stages. Since *T*_eye_ is relatively well correlated to core temperature in chicken (Cândido et al. 2020), budgerigars (*Melopsittacus undulates*; Ikkatai and Watanabe 2015) and wild red-fotted boodies (*Sula sula*; Gauchet et al. 2022), our results suggest that core body temperature increased during the early stage of moult (+1.61°C of *T*_eye_ between old plumage and 25% stage). This idea is supported by an increase of *ca*. 0.8°C in core temperature during moult in Yellow-eyed penguin (*Megadyptes antipodes*; Farner 1958). Yet, blood flow to the eyes, impacting *T*_eye_, has been also shown to play a role in reducing brain temperature in pigeons (*Columba livia*; Pinshow et al. 1982), suggested as heat sink in ostriches (*Struthio camelus*; Fuller et al. 2003) or to maintain/enhance visual acuity during stress (Winder et al. 2020). Unfortunately, without core body temperature measurements, it is not possible here to formally assess if the increased heat dissipation through thermal windows we observed was sufficient to maintain core body temperature at stable levels, or if the observed increase in *T*_eye_ could reflect the inability to maintain core body temperature at stable levels.

Interestingly, the thermal challenge to dissipate heat described here at thermoneutrality seems likely to be specific to penguins, due to the accumulation of this double insulation. Indeed, most studies in birds measured the opposite pattern with an increase of 30 to 60% in thermal conductance during moult (Lustick 1970; Dietz et al. 1992), inducing for instance a rise of the lower critical temperature in Long-eared Owl (*Asio otus*; Wijnandts 1984). In seals, moult represents the renewal not only of the hair but also of the skin (Ling 1968; 1972). With no accumulation of old and new skin, individuals show an increase of thermoregulation cost due to higher heat loss (Paterson et al. 2012). In Antarctica Weddell seals (*Leptonychotes weddellii*), Walcott et al. (2020) measured that the energetic costs of thermoregulation doubled during moulting period, supported by an increase of 25% of heat loss in early moulting stages. Importantly, no thermal windows were detected in this study suggesting that individuals were unlikely to overheat (Walcott et al. 2020). Similar results were obtained in southern elephant seals (*Mirounga leonina*) with an increase of 1.8 × resting metabolic rate during moult with body surface temperature decreasing throughout the moult (Paterson et al. 2022), the latter suggesting an energy-saving strategy that seems opposite to the results obtained in this study in Gentoo penguins maintained within their thermoneutral zone. Moreover, elephant seals also showed aggregation behavior only during moult on land suggesting a strategy to reduce heat loss and minimize energy costs (Chaise et al. 2019). In contrast, penguins have not been shown to exhibit specific aggregation behavior during moulting to the best of our knowledge.

Surface temperatures of new plumage stage did not reach the initial temperature measured in old plumage and they were either lower (*T*_bill_, *T*_eye_ and *T*_flipper_) or similar (*T*_foot_; *T*_trunk_) at the end of the moult. Moult could have started before a visible change in plumage (i.e. pop-corn stage). In this case, the old plumage stage (M1) could already correspond to an early stage of moult (i.e. growth of new feathers below the skin, Fig. 1), with a higher metabolism, rather than a pre-moult stage as initially considered in this study. Secondly, the new plumage stage (M7) corresponds here to the end of the old feather loss and potentially not to the end of the new feather growth. The immature new feathers could be less insulated than the full-length feathers, allowing heat loss without a specific need to maintain blood flow to thermal windows. Finally, since moulting progression is also associated with the progression into a more advanced fasting stage, it is possible that individuals reaching the new plumage stage could use peripheral vasoconstriction as an energy-saving strategy (Kooyman et al. 1976; Ponganis et al. 2001, 2003; Tattersall et al. 2016) which could also explain the decrease we observed in *T*_bill_, *T*_eye_ and *T*_flipper_.

Our study showed that under nearly constant environmental conditions, moulting Gentoo penguins increase the surface temperature of poorly insulated regions (thermal windows) (Fig. 2 and 3) to dissipate extra heat. In the wild, the thermal challenge of carrying two feather layers could be enhanced by the solar radiation and the increase of air temperature in the current context of global warming (Ainley et al. 2010; Gorodetskaya et al. 2022). Further studies are needed to better understand the operative temperature (Bakken 1976) experienced by wild birds, especially in polar marine species showing adaptations to reduce heat loss while foraging in cold water.

## Data availability statement

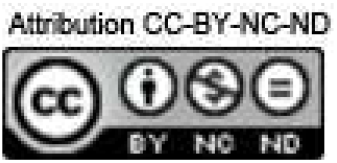

Anyone can share this material, provided it remains unaltered in any way, this is not done for commercial purposes, and the original authors are credited and cited. https://figshare.com/s/c64975557480cc8709f3

## Acknowledgments

We are grateful to Oceanopolis© for providing logistical support and giving us the access the animals. We would like to thank Christine Dumas, Alexiane Corcuff, Maxence Leroy, Mélanie Robert and Agathe Lefranc for their help during experiment. We sincerely thank Dominic McCafferty for his valuable comments improving a previous version of the manuscript. AL was supported by ISblue project, Interdisciplinary graduate school for the blue planet (ANR-17-EURE-0015) and co-funded by a grant from the French government under the program “Investissements d’Avenir” embedded in France 2030. AS was financially supported by the CNRS and the IdEx Université de Strasbourg.

